# Optimal dynamic incentive scheduling for Hawk-Dove evolutionary games

**DOI:** 10.1101/2021.08.15.456406

**Authors:** K. Stuckey, R. Dua, Y. Ma, J. Parker, P.K. Newton

## Abstract

The Hawk-Dove mathematical game offers a paradigm of the trade-offs associated with aggressive and passive behaviors. When two (or more) populations of players (animals, insect populations, countries in military conflict, economic competitors, microbial communities, populations of co-evolving tumor cells, or reinforcement learners adopting different strategies) compete, their success or failure can be measured by their frequency in the population (successful behavior is reinforced, unsuccessful behavior is not), and the system is governed by the replicator dynamical system. We develop a time-dependent optimal-adaptive control theory for this nonlinear dynamical system in which the payoffs of the Hawk-Dove payoff matrix are dynamically altered (dynamic incentives) to produce (bang-bang) control schedules that (i) maximize the aggressive population at the end of time *T*, and (ii) minimize the aggressive population at the end of time *T*. These two distinct time-dependent strategies produce upper and lower bounds on the outcomes from all strategies since they represent two extremizers of the cost function using the Pontryagin maximum (minimum) principle. We extend the results forward to times *nT* (*n* = 1, …, 5) in an adaptive way that uses the optimal value at the end of time *nT* to produce the new schedule for time (*n* + 1)*T*. Two special schedules and initial conditions are identified that produce absolute maximizers and minimizers over an arbitrary number of cycles for 0 ≤ *T* ≤ 3. For *T* > 3, our optimum schedules can drive either population to extinction or fixation. The method described can be used to produce optimal dynamic incentive schedules for many different applications in which the 2 × 2 replicator dynamics is used as a governing model.

## I. INTRODUCTION

The Hawk-Dove game (aka Chicken or Snowdrift game) is a game-theoretic paradigm for studying the conflict between players (or populations of players) who use two opposing strategies: aggressive (Hawks) and passive (Doves). One way of framing the conflict is to consider competition in the animal world where two different species compete for a limited resource [1–4]. With no Hawks in the population, Doves will share the resources and avoid conflict. With no Doves, the Hawks will fight with each other for resources, taking the risk of injury or death. If Hawks are present in large enough numbers, the Doves will flee without fighting. A sufficient fraction of Doves, on the other hand, can cooperate and expel the Hawks from the population thereby protecting the resource [5]. The challenge is to find conditions for stable co-existence of the two opposing populations. In the context of military conflicts, the game is framed as the game of chicken, thought of as a situation in which two drivers head towards each other in a single lane trying not to be the first to swerve away (Doves), each mindful of the fact that if neither swerves (Hawks), both will die. Key to this game is that the cost of losing is greater than the value of winning. Versions of this (static) game have been analyzed and used extensively in political science communities to study strategies associated with the problem of nuclear brinkmanship [6]. In this set-up, the payoffs are fixed, and the interactions unfold based on the cost-benefit balance determined by these payoffs.

In a more complicated setting, one might want to measure repeated interactions in populations of competitors, 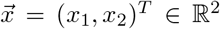, where winning and losing is reinforced by the relative frequencies of the two competing populations (frequency dependent selection as in Darwinian evolution). For this, the replicator dynamical system is commonly used [7–9]:

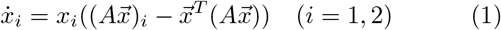

with *x*_1_ + *x*_2_ = 1, 0 ≤ *x*_1_ ≤ 1, 0 ≤ *x*_2_ ≤ 1, where each variable has the interpretation of frequency in the population or the alternative interpretation as a probability of picking a member of one of the two subgroups randomly. It is useful to also think of the variables 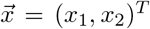 as strategies (heritable traits) that evolve, with the most successful strategy dominating, as in the context of Darwinian evolution [4] by natural selection. Here, *A* is the 2 × 2 payoff matrix, 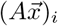 is the fitness of population *i*, and 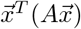 is the average fitness of both populations, so *x_i_* in (1) drives growth if the population *i* is above the average and decay if it is below the average. The fitness functions in (1) are said to be population dependent (selection pressure is imposed by the mix of population frequencies) and determine growth or decay of each subpopulation. Because of this, these equations are also used extensively in the reinforcement learning community where success begets success and failure leads to a downward spiral of frequency in the population [10].

Using the standard Hawk-Dove payoff matrix [5]:

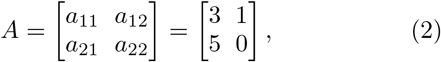

where the population *x*_1_ are the Hawks (aggressive), and *x*_2_ are the Doves (passive), the strict Nash equilibrium, 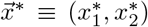 is the mixed state 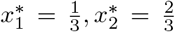 since 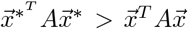 for all 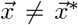. This implies that the mixed state is also an evolutionary stable state (ESS) of the replicator system (1) as discussed in [11]. It is also useful to uncouple the two variables in (1) and write a single equation for the aggressor population frequency *x*_1_:

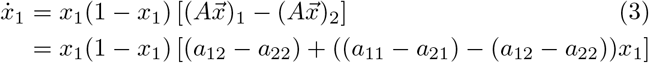

Note also that a single equation for the passive population *x*_2_ is easily obtained using the change of variable *x*_1_ = 1 − *x*_2_ in eqn (3).

The question we address in the paper is whether it is possible to alter the entries in the payoff matrix *A* in a time-dependent fashion (dynamic incentives) in order to optimally achieve some pre-determined goal (such as minimizing aggression) at the end of fixed time *T*? Dynamically altering the entries of a payoff matrix in an evolutionary game setting has only recently been studied by coupling the entries, for example, to a system that represents an external environment [12, 13]. In the context of nuclear brinkmanship, is it possible to alter the payoff incentives dynamically in order to achieve a goal [6] that would not be achievable with fixed payoffs? Is it possible to offer dynamic economic incentives that optimize some desired outcome across a population of participants [14, 15]? Can one optimally design time-dependent incentive schedules of rewards/punishments to compel groups of people to get vaccinated [16]? For co-evolving microbial populations, is it possible to dynamically schedule selective antibiotic agents in order to *steer* the evolutionary trajectory in an advantageous direction [17, 18], or even reverse antibiotic resistance, or in the context of scheduling chemotherapy treatments, is it possible to design schedules optimally that make best use of the chemotherapy agents administered in order to delay chemotherapeutic resistance [8, 9, 19–21]? Control theory is increasingly being used in a wide range of biological applications [21–27] but to date, has not been systematically implemented in the context of evolutionary games as far as we know, aside from [8, 9, 21, 28].

One evolutionary context where an apparent Hawk-Dove scenario may require attainment of a quasi-stable equilibrium condition is during the evolution of symbiotic relationships in which one partner is aggressive or predatory. For example, hostile colonies of eusocial insects, such as ants and termites, are plagued by a diversity of solitary arthropods that have evolved to infiltrate the social system and parasitize the nest [30, 31]. The majority of such parasitic species evolved from free-living ancestors without any behavioral specialization [32, 33]. It follows that the initial steps in establishing the symbiosis were contingent on these free-living species (the Doves) entering into equilibrium with their aggressive eusocial hosts (the Hawks). This equilibrium, once attained, may have provided an essential, permissive stepping stone to evolving the essential adaptive traits—such as social behaviors and pheromonal mimicry—that facilitate social parasitism [32].

To address these and related types of settings, we develop a mathematical framework to determine time-dependent incentive schedules for altering the payoff entries of a Hawk-Dove evolutionary game in such a way as to (i) maximize aggression at the end of time *T*, and (ii) minimize aggression at the end of time *T*. By considering the bang-bang schedules that produce these upper and lower bounds on the competing frequencies, we can conclude that any alternative payoff schedule will produce a result that lies somewhere between the two bounds as each are extremizers of a cost function associated with the Pontryagin maximum (minimum) principle. We then extend the time-period to time *nT* (*n* = 1, …, 5) by using an adaptive control method that adjusts the schedule in the (*n* + 1)*^st^* window based on the ending frequency values from the *n^th^* window. The schedules produced drive aggression down to an absolute minimum 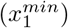, or drive it up to an absolute maximium 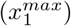, which are functions of the cycle-time *T*. These values provide absolute lower and upper bounds on opposing behavior strategies in an evolutionary setting.

## II. OPTIMAL CONTROL THEORY FOR THE REPLICATOR DYNAMICAL SYSTEM

To implement an optimal dynamic incentive strategy, we consider the system:

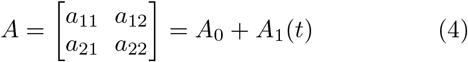

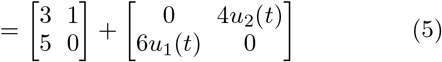

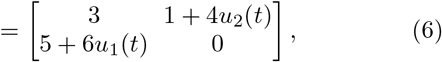

where *A*_1_(*t*) represents our control with entries in the off-diagonal terms, and *A*_0_ is the baseline Hawk-Dove payoff matrix. The time-dependent functions 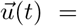 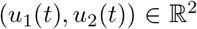 are bounded above and below, − 1 ≤ *u*_1_(*t*) ≤ 1, − 1 ≤ *u*_2_(*t*) ≤ 1 and have a range (− 3 ≤ *a*_12_ ≤ 5; − 1 ≤ *a*_21_ ≤ 11) that allows us to traverse the plane along any path depicted in red in figure 1, starting in the Hawk-Dove zone in the uncontrolled (*u*_1_ = 0; *u*_2_ = 0) case which is shown in figure 2 in the phase plane (a) and the frequency plane (b). The ESS for the uncontrolled case is *x*_1_ = 1/3. The control path chosen, and the time parametrization 0 ≤ *t* ≤ *T* determines both the sequence of games being played as well as the switching times (the times at which the path crosses over from one region to the next) between games. We denote the total control output 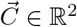:

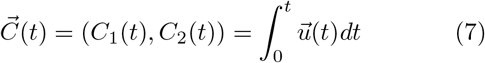

with total output delivered in time *t*, then:

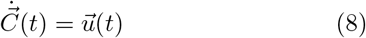

and:

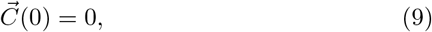

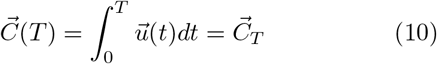

where *T* denotes a final time in which we implement the control over one cycle. We consider 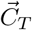 as a constraint on the optimization problem, with 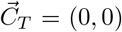, and our goal is to first find schedules that minimize and maximize aggression (*x*_1_) at the end of one cycle *t* = *T* subject to this constraint. For the uncontrolled case, we know *x*_1_ → 1/3 as *t* → ∞ and we compare the controlled cases with the uncontrolled case, both satisfying the constraint. Also notice that the linear growth rate in (3) is (*a*_12_ − *a*_22_) = 1 − 0 = 1, so we scale *T* the same way in our computations, as *T* = 1. We then perform the optimization adaptively over multiple cycles *nT* using the end value of cycle *nT* as the initial condition to compute the optimal schedule for the (*n* + 1)*^st^* cycle. Using this method, we are able to identify absolute maximizers and minimizers as a function of the cycle time *T*.

**FIG. 1.**
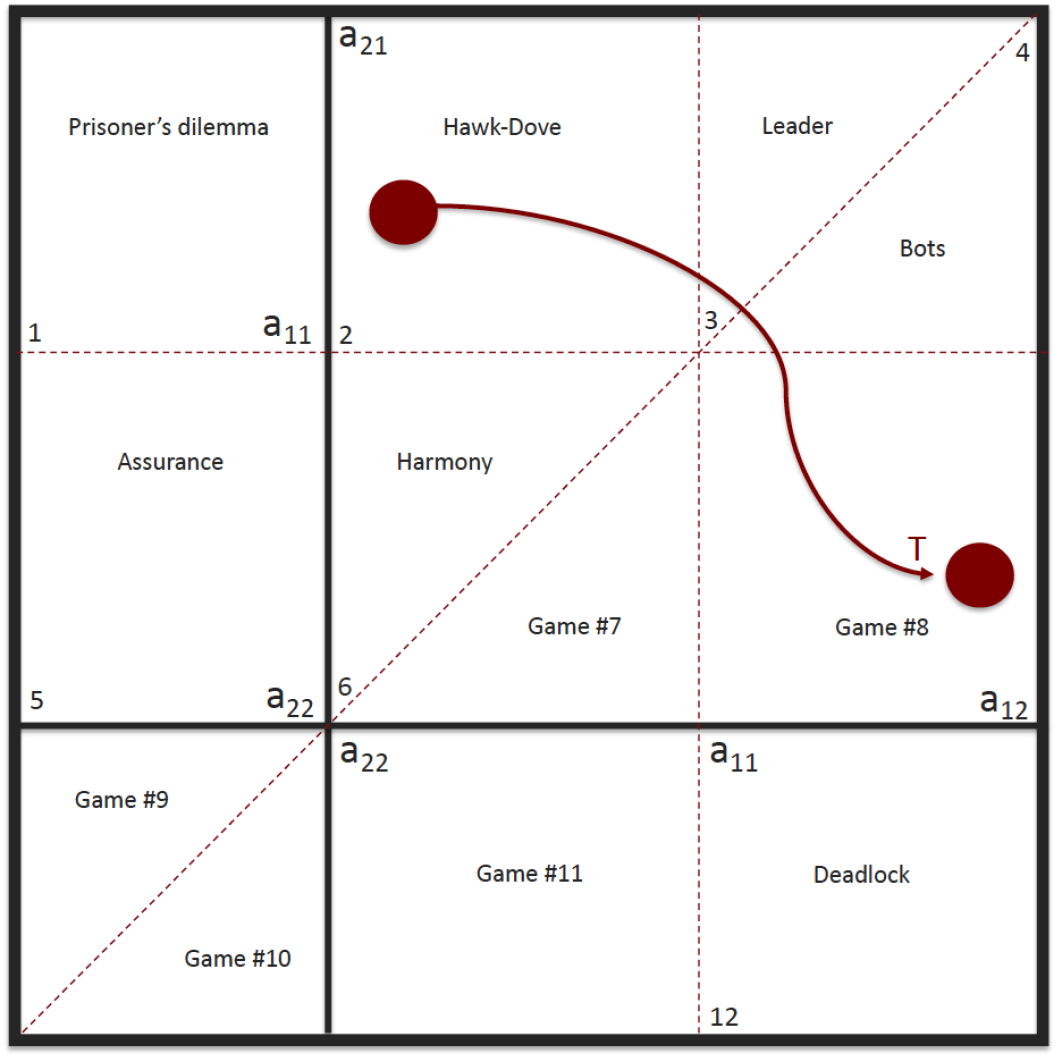
Twelve regions in the (*a*_12_, *a*_21_) plane [29] define which game is being played. We choose *a*_22_ at the origin (without loss). Starting at *t* = 0 in the Hawk-Dove square, what are the paths to travel that minimize and maximize aggression at time *t* = *T*?

**FIG. 2.**
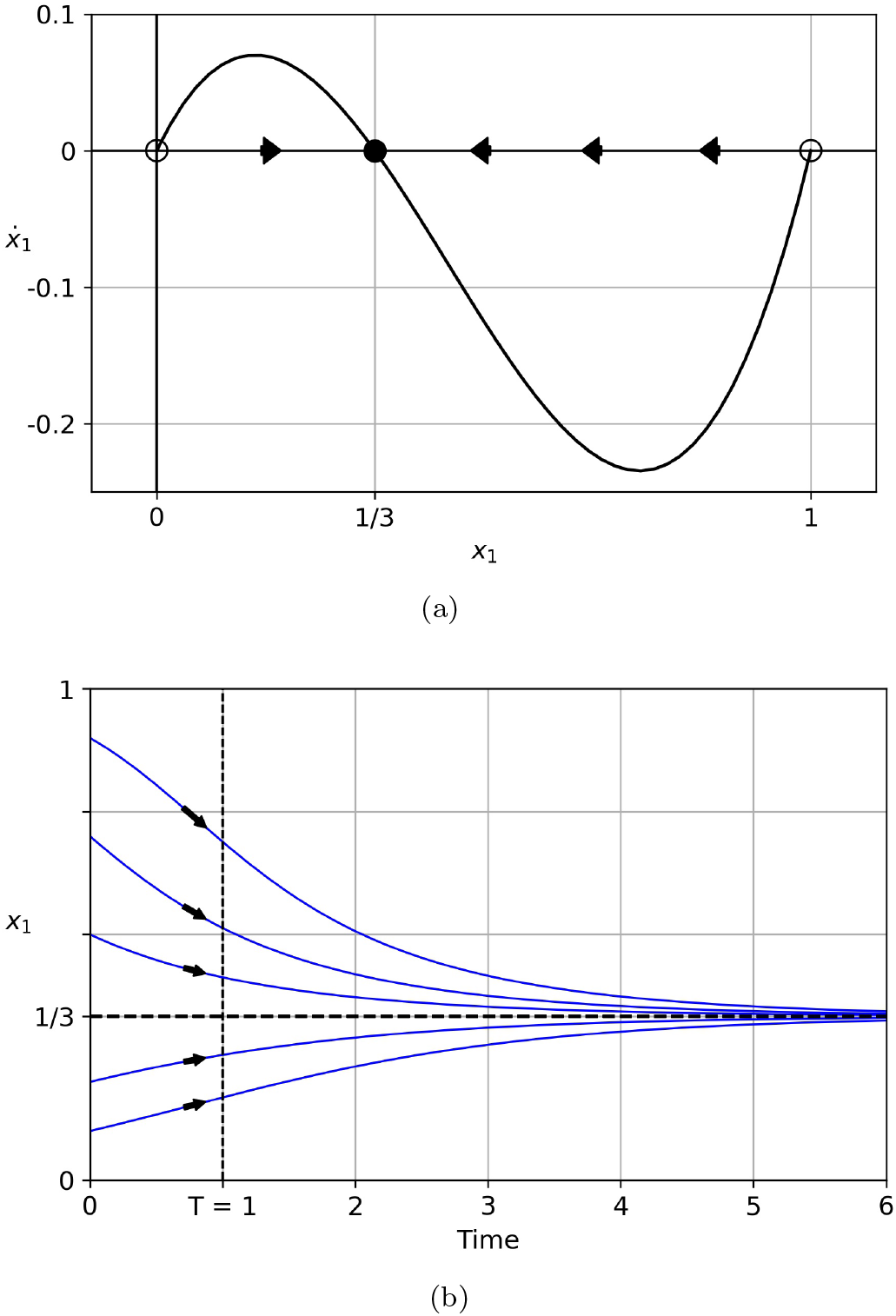
Dynamics of the uncontrolled (*u*1 = 0, *u*2 = 0) Hawk-Dove evolutionary game. (a) Phase portrait associated with the aggressor population *x*_1_. Both Hawk and Dove dominance (*x*_1_ = 1, 0) are unstable fixed points, while the mixed state *x*_1_ = 1/3 is the evolutionarily stable strategy (ESS); (b) Hawk dynamics for various initial conditions. *T* = 1 is the end of one control cycle and also the linear growth rate of the Hawk-Dove system.

### A. Optimal control formulation

A standard form for implementing the Pontryagin maximum (minimum) principle with boundary value constraints is:

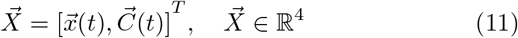

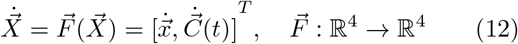

where we would like to minimize or maximize a general cost function:

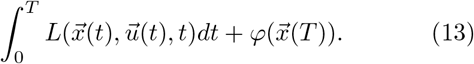

Since the method is standard, we will just briefly describe the basic framework and refer readers to [34–38] for more details on how to implement the approach. Following [37] in particular (see page 62 Theorem 4.2.1), we construct the control theory Hamiltonian:

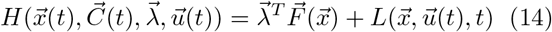

where 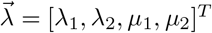 are the co-state functions (i.e. momenta) associated with 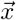 and 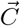 respectively. Assuming that 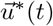 is the optimal control for this problem, with corresponding trajectory 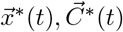, the canonical equations satisfy:

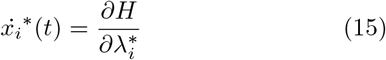

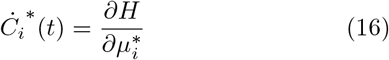

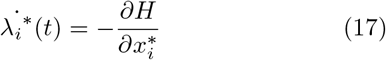

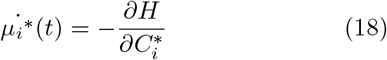

where *i* = (1, 2). The corresponding boundary conditions are:

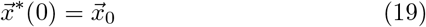

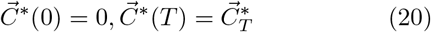

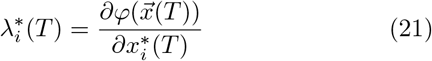

Then, at any point in time, the optimal control 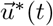 will minimize the control theory Hamiltonian:

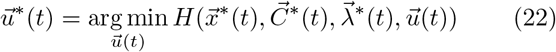

The optimization problem becomes a two-point boundary value problem (using (19)–(21)) with unknowns 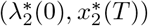 whose solution gives rise to the optimal trajectory 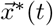 (from (15)) and the corresponding control 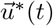 that produces it [34–37]. We choose our cost function (13):

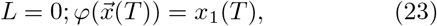

and we solve this problem by standard numerical shooting type methods [37].

## III. RESULTS

In this section we show the results of the adaptive optimal control method to minimize and maximize aggression at time *T* = 1, and then further at the end of multiple cycles *t* = *nT*. Figure 3(a)–(i) shows the maximizing (blue) and minimizing (red) trajectories for nine initial conditions. The corresponding bang-bang schedules that produce these trajectories are also shown in each case. It is straighforward to prove that the optimal schedules must be bang-bang since the controllers are linear in the governing equations. In each case, we show the uncontrolled (dashed curve) Hawk-Dove trajectory, which ends in between the maximizer and minimizer as expected.

**FIG. 3.**
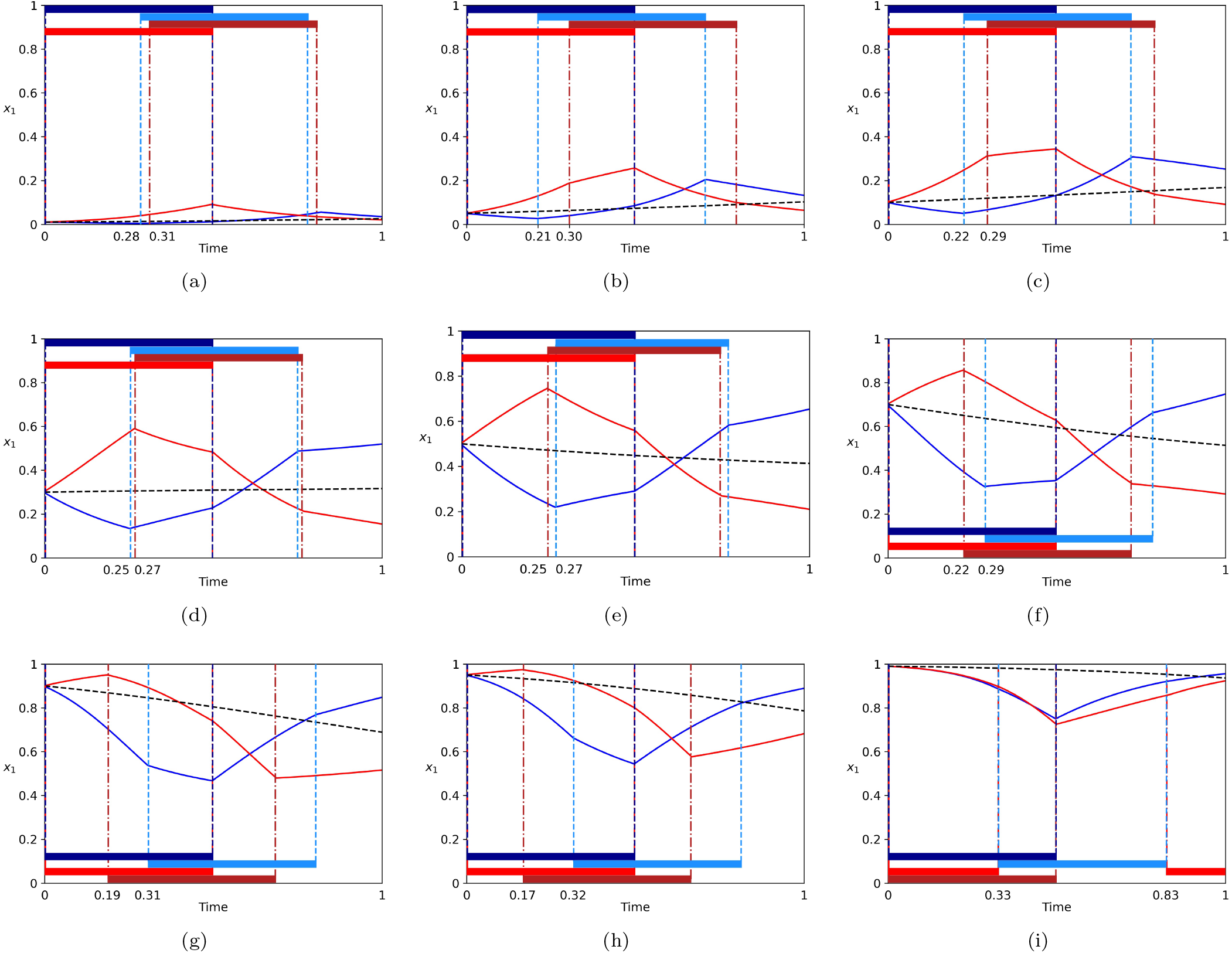
Maximizing (blue) and minimizing (red) trajectories for nine initial conditions. Dashed curve shows the uncontrolled Hawk-Dove trajectory which lands in between the max and min at *T* = 1. Dark (light) blue bar shows *u*_1_ = 1 (*u*_2_ = 1), white bar shows *u*_1_ = − 1 (*u*_2_ = − 1) associated with the maximizing control schedule; Dark (light) red bar shows *u*_1_ = 1 (*u*_2_ = 1), white bar shows *u*_1_ = − 1 (*u*2 = − 1) associated with the minimizing control schedule. All schedules are bang-bang. (a) *x*_1_(0) = 0.01; (b) *x*_1_(0) = 0.05; (c) *x*_1_(0) = 0.1; (d) *x*_1_(0) = 0.3; (e) *x*_1_(0) = 0.5; (f) *x*_1_(0) = 0.7; (g) *x*_1_(0) = 0.9; (h) *x*_1_(0) = 0.95; (i) *x*_1_(0) = 0.99.

Figure 4 shows the maximizing (blue) and minimizing trajectories over *n* = 5 cycles. We obtain these adaptively, using the endpoint from the *n^th^* cycle to compute the optimal schedule for the following (*n* + 1)*^st^* cycle. Two special initial conditions are shown in figure 5. For *x*_1_(0) = 0.08, the minimizing (red) trajectory shown in figure 5(a) ends at *x*_1_(1) = 0.08, hence is periodic. This value (and corresponding schedule) corresponds to an absolute minimizer 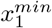 for aggression *x*_1_. By contrast, for *x*_1_(0) = 0.79 shown in figure 5(b), the maximizing (blue) trajectory ends at *x*_1_(1) = 0.79, hence is periodic. This value (and the corresponding schedule) corresponds to an absolute maximizer 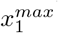 for aggression. These two special initial conditions are shown in figure 6 over *n* = 5 cycles confirming the periodicity of the minimizing trajectory (red) in figure 6(a) and the periodicity of the maximizing (blue) trajectory in figure 6(b). The sequence of games that the system cycles through to achieve the minimizing sequence is shown in figure 7, while the maximizing sequence is shown in figure 8. These are obtained from eqn (3) and the four equations:

1. *u*_1_ = 1; *u*_2_ = 1: 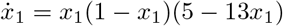
2. *u*_1_ = 1; *u*_2_ = − 1: 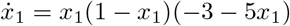
3. *u*_1_ = − 1; *u*_2_ = 1: 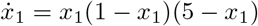
4. *u*_1_ = − 1; *u*_2_ = − 1: 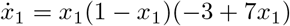.

**FIG. 4.**
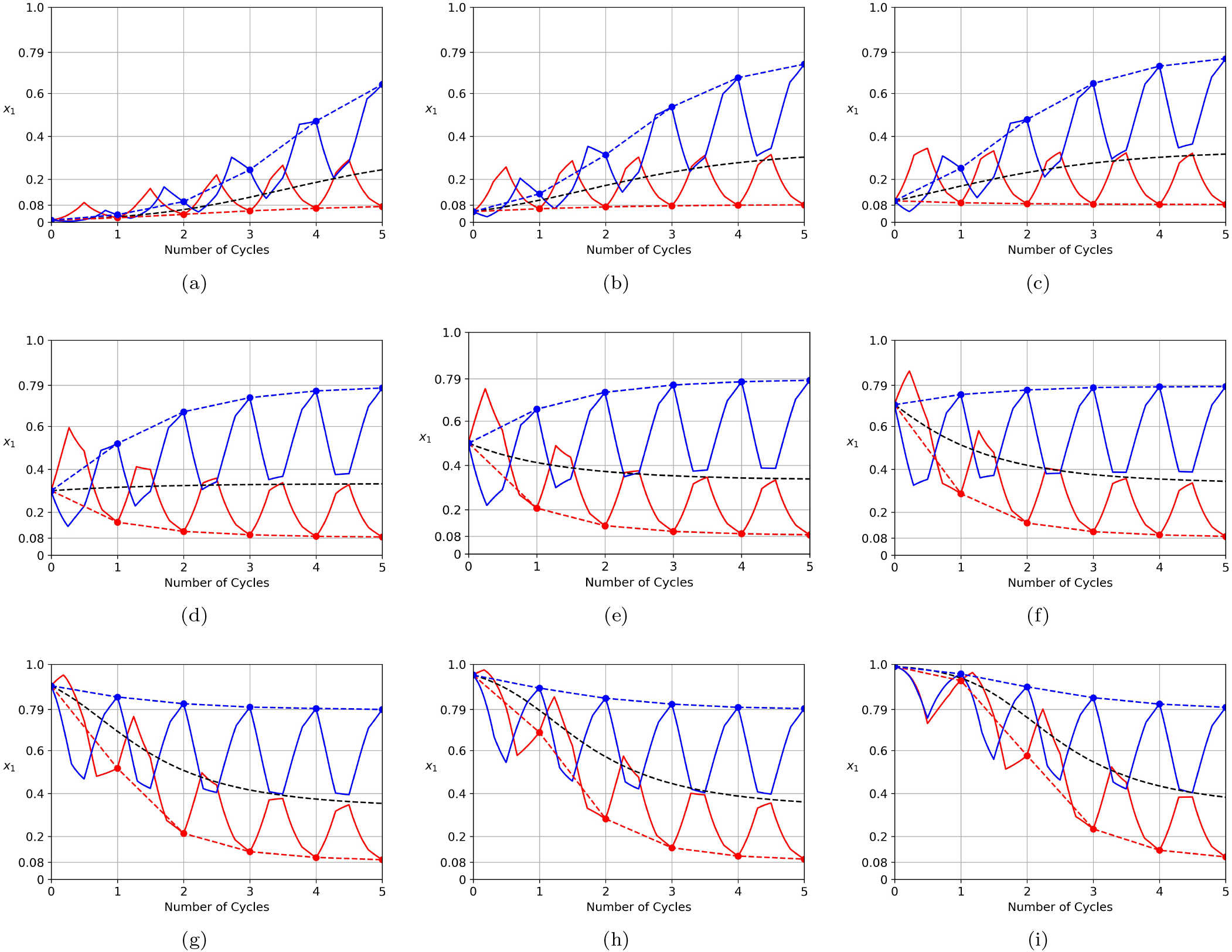
Maximizing (blue) and minimizing (red) trajectories for nine initial conditions over *n* = 5 cycles, with dashed blue/red curves joining the end values after each cycle. The adaptive schedule for the (*n* + 1)*^st^* cycle is calculated based on the endpoint of the *n^th^* cycle. Black dashed curve shows the uncontrolled Hawk-Dove trajectory. (a) *x*_1_(0) = 0.01; (b) *x*_1_(0) = 0.05; (c) *x*_1_(0) = 0.1; (d) *x*_1_(0) = 0.3; (e) *x*_1_(0) = 0.5; (f) *x*_1_(0) = 0.7; (g) *x*_1_(0) = 0.9; (h) *x*_1_(0) = 0.95; (i) *x*_1_(0) = 0.99.

**FIG. 5.**
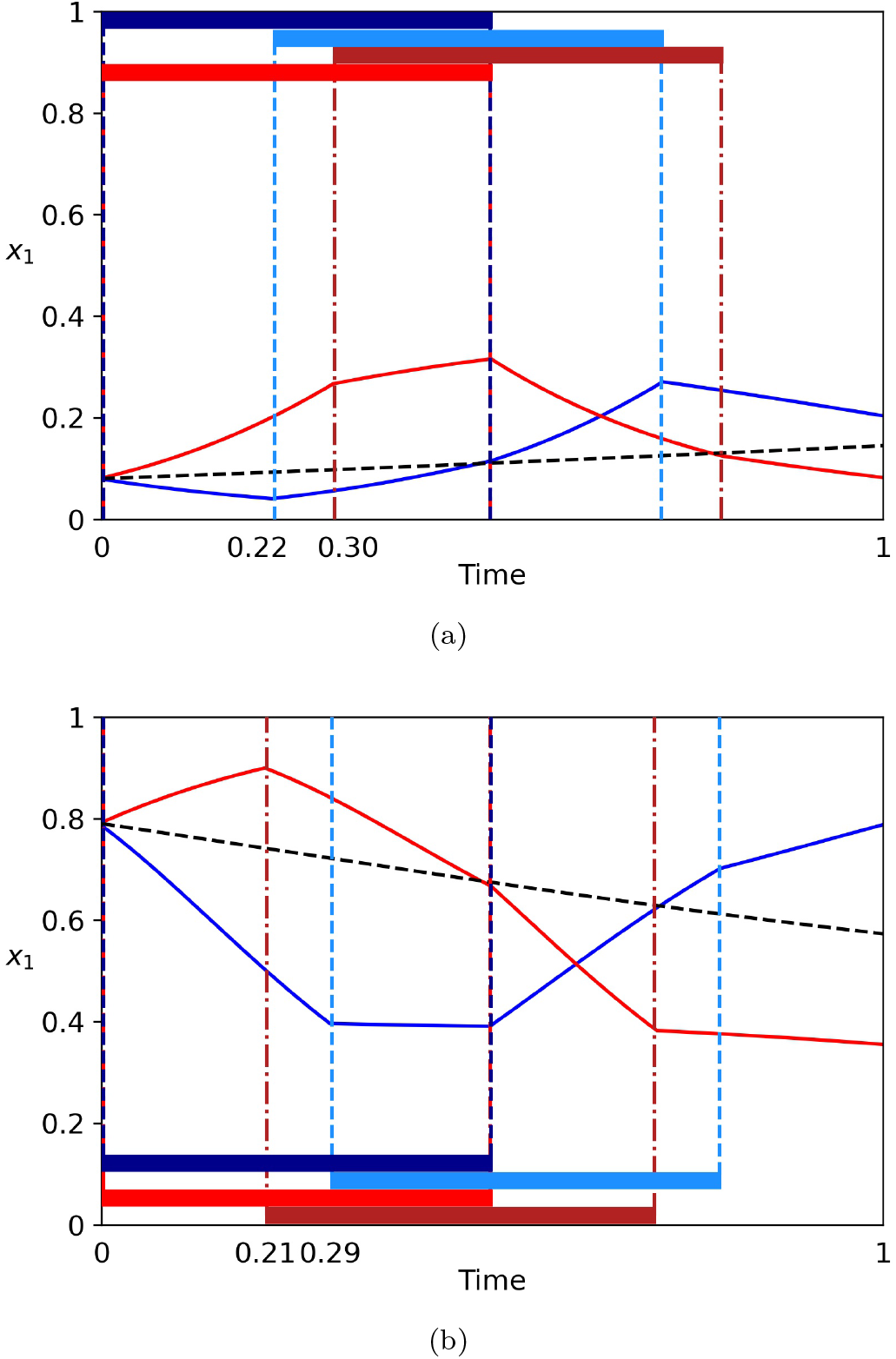
Maximizing (blue) and minimizing (red) trajectories for the two special initial conditions *x*_1_(0) = 0.08, 0.79. For the larger initial condition, the maximizing schedule (blue) produces an absolute maximum 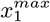 for *T* = 1. For the smaller initial condition, the minimizing schedule (red) produces an absolute minimum 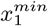 for *T* = 1. Dashed curve shows the uncontrolled Hawk-Dove trajectory. Dark (light) blue bar shows *u*_1_ = 1 (*u*_2_ = 1), white bar shows *u*_1_ = − 1 (*u*2 = − 1) associated with the maximizing control schedule. (a) *x*_1_(0) = 0.08; (b) *x*_1_(0) = 0.79.

**FIG. 6.**
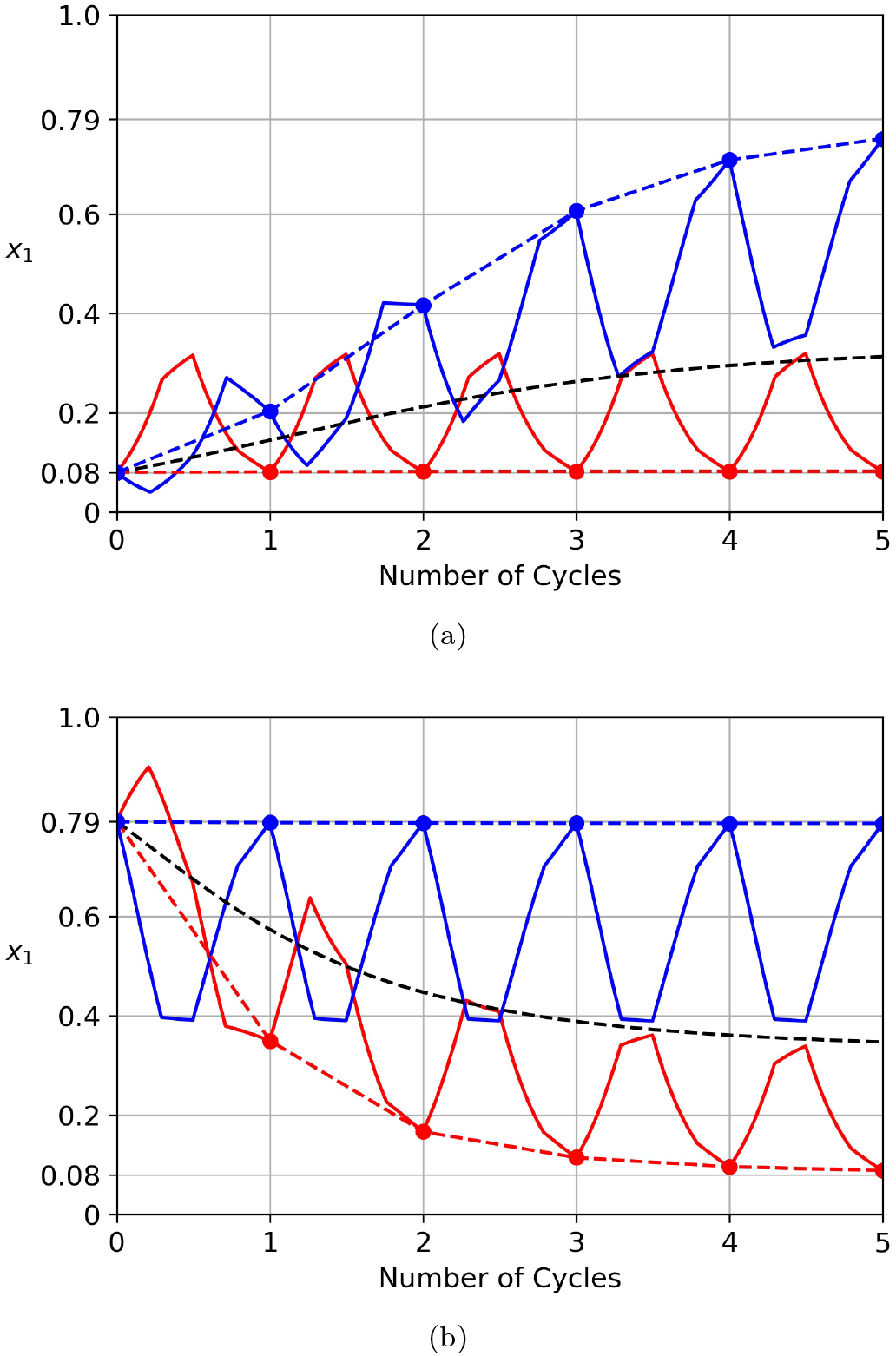
Maximizing (blue) and minimizing (red) trajectories for the two special initial conditions *x*_1_(0) = 0.08, 0.79 over *n* = 5 cycles. Notice that the minimizing trajectory (red) shown in (a) exactly repeats for each cycle (*x*_1_(0) = *x*_1_(*T*)) since the schedule is an absolute minimizer, while the maximizing trajectory (blue) shown in (b) exactly repeats for each cycle (*x*_1_(0) = *x*_1_(*T*)) since the schedule is an absolute maximizer. Dashed curve shows the uncontrolled Hawk-Dove trajectory. (a) *x*_1_(0) = 0.08. Dashed red horizontal line indicates an absolute minimizer at 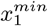; (b) *x*_1_(0) = 0.079. Dashed blue horizontal line indicates an absolute maximizer at 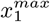.

**FIG. 7.**
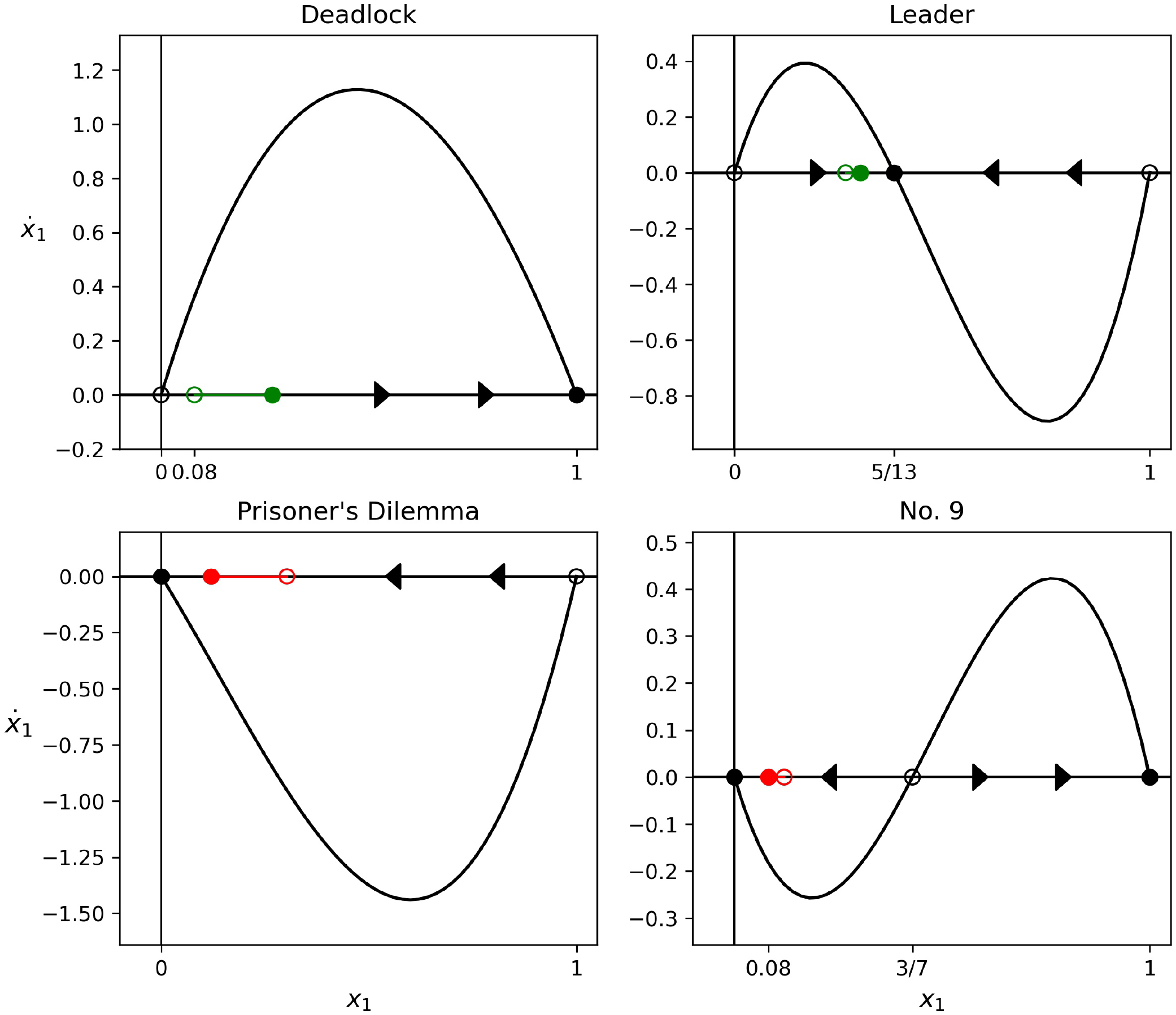
Minimizing sequence of four games (Deadlock; Leader; Prisoner’s Dilemma; Game 9) associated with initial condition *x*_1_(0) = 0.08 that produces the absolute minimizer. Red line marks the distance decreased from initial point (unfilled red dot) to final point (filled red dot). Green line marks the distance increased from initial point (unfilled green dot) to final point (filled green dot).

**FIG. 8.**
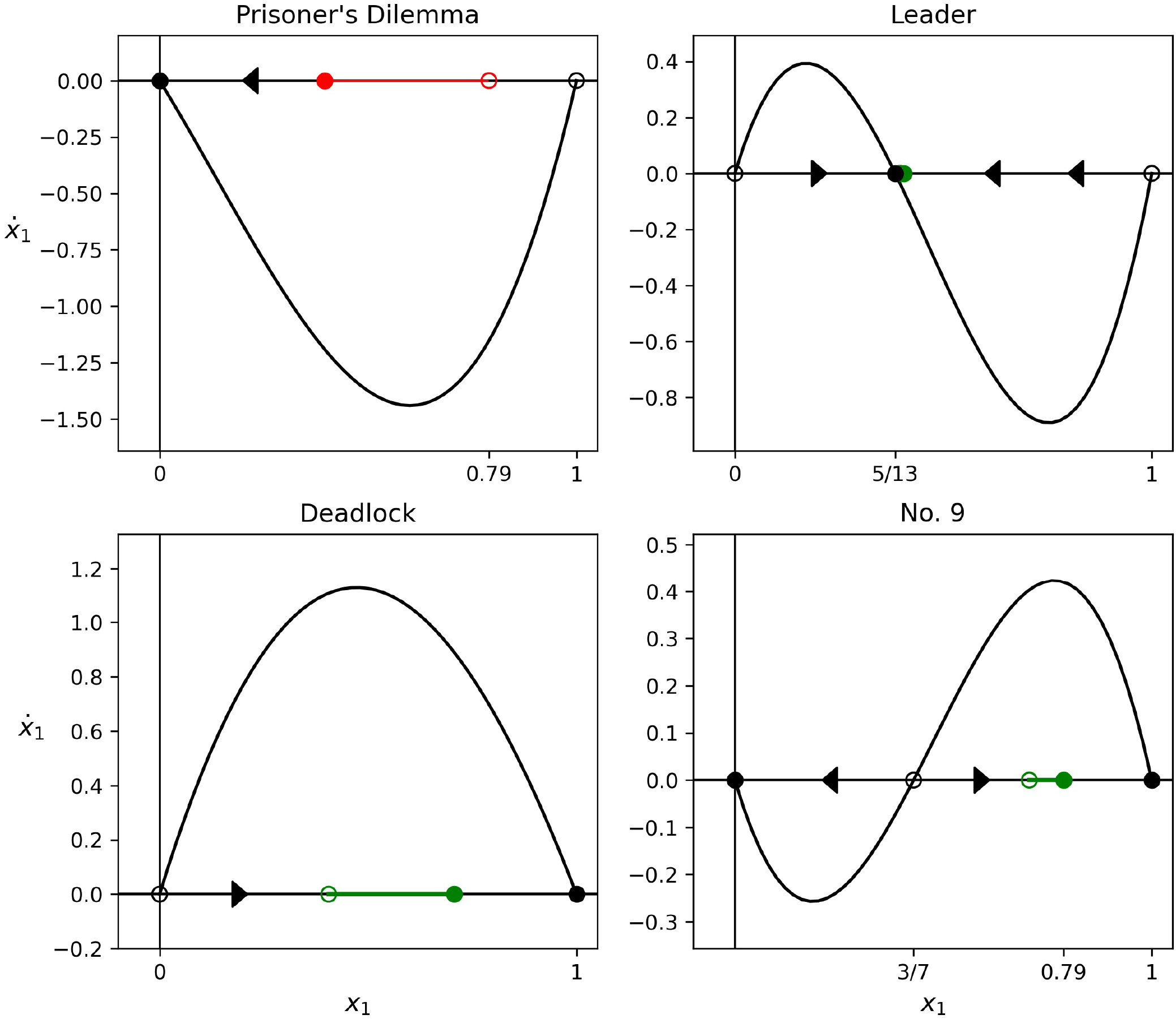
Maximizing sequence of four games (Prisoner’s Dilemma; Leader; Deadlock; Game 9) associated with initial condition *x*_1_(0) = 0.79 that produces the absolute maximizer. Red line marks the distance decreased from initial point (unfilled red dot) to final point (filled red dot). Green line marks the distance increased from initial point (unfilled green dot) to final point (filled green dot).

In figure 9 we show the minimizing values and maximizing values of *x*_1_(*T*) vs. *x*_1_(0) through the full range 0 ≤ *x*_1_(0) ≤ 1. Notice that at the endpoints, the two values converge (since for the linear system, the schedule does not matter, only the total 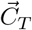). Figure 9(b) shows the ratio *x*_1_(*T*)/*x*_1_(0) − 1 (percentage increase or decrease) vs. initial condition *x*_1_(0) through the full range 0 ≤ *x*_1_(0) ≤ 1. When the maximizing (blue) curve crosses *x*_1_(*T*)/*x*_1_(0) − 1 = 0, (i.e. *x*_1_(0) = *x*_1_(*T*)) an absolute maximizer is achieved (for 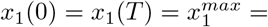 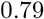), while when the minimizing (red) curve crosses *x*_1_(*T*)/*x*_1_(0) − 1 = 0, an absolute minimizer is achieved (for 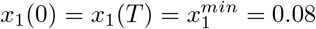). In figure 10 we show how 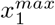 and 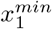 depend on the cycle-time *T*. Interestingly, as *T* → 0, 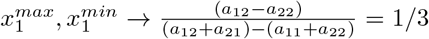 which is the ESS for the uncontrolled Hawk-Dove system. For 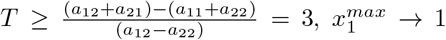 and 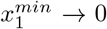 showing that for large enough cycle times we can drive either of the sub-populations to extinction or to fixation.

**FIG. 9.**
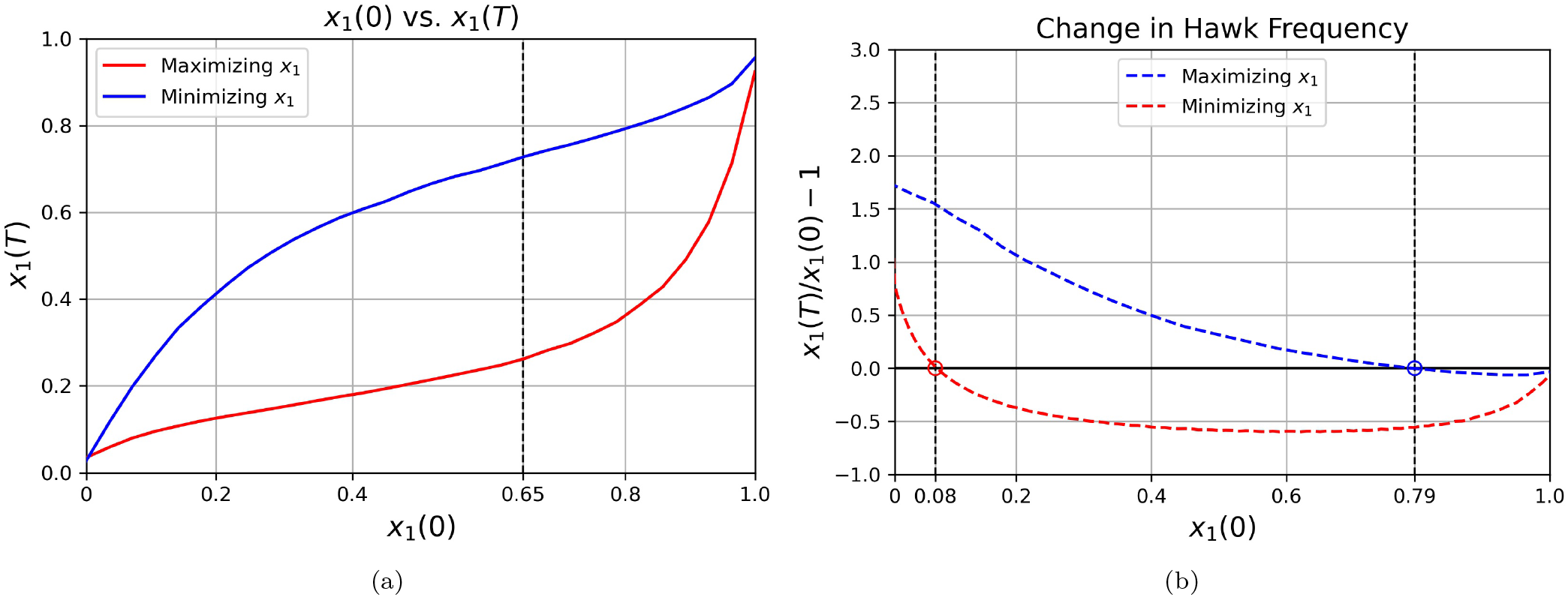
(a) Hawk initial condition *x*_1_(0) versus Hawk frequency at final time *x*_1_(*T*) for maximizing (blue) and minimizing (red) schedules. Vertical dashed line at *x*_1_(0) = 0.65 marks the maximum difference between the minimizer and the maximizer; (b) Change in Hawk frequency as a function of initial condition. Points above the line *x*_1_(*T*)/*x*_1_(0) − 1 = 0 represent an increase over time and points below this line represent a decrease over time. The two intersection points *x*_1_(0) = 0.08 and *x*_1_(0) = 0.79 mark the absolute minimizer and maximizer initial conditions for *T* = 1.

**FIG. 10.**
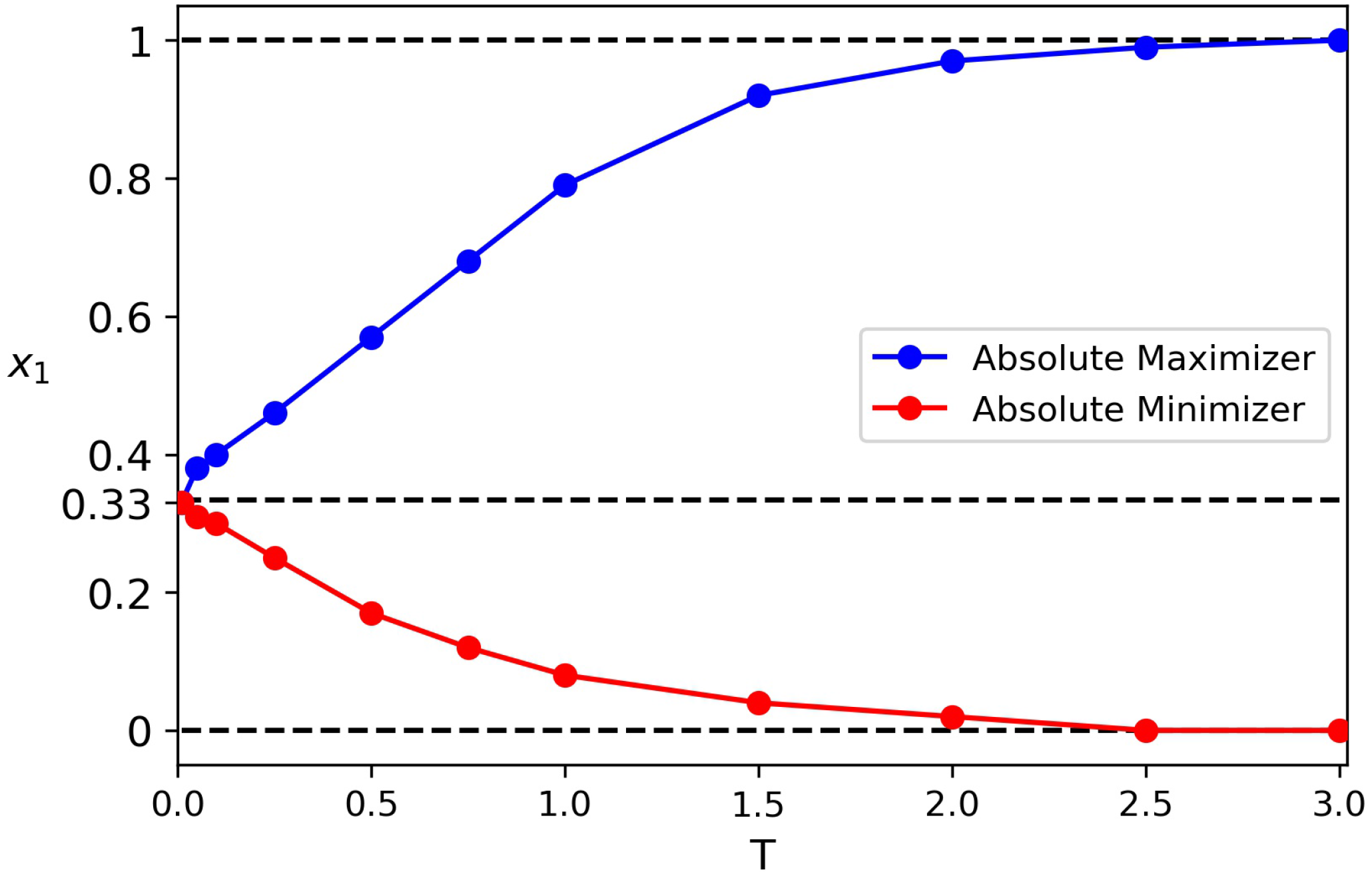
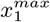 (blue) and 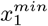 (red) as a function of cycle-time *T*. Dashed horizontal line at *x*_1_ = 1/3 is the ESS for the uncontrolled Hawk-Dove system where the two curves meet as *T* → 0.

## IV. DISCUSSION

Our goal in this manuscript is to lay out the general mathematical framework for determining optimal dynamic incentive schedules (time-dependent payoff schedules) that maximize/minimize certain behaviors in an evolutionary game theory setting using the 2 × 2 replicator. dynamical system with a Hawk-Dove payoff matrix as our baseline. By changing the payoff entries in a time-dependent manner, subject to constraints, we are altering the payoff-reward structure of the Hawk-Dove interaction as the populations evolve, which is equivalent to selecting a sequence of 2 × 2 evolutionary games in such a way that an optimum is achieved after a fixed passage of time. The determination of these schedules requires a balance between the timescale on which the payoffs change, and the timescale of the underlying replicator dynamical system in such a way that the Pontryagin maximum/minimum principle is satisfied.

As mentioned earlier, there are many settings in which dynamic payoffs can be used to achieve a certain out-come (developing chemotherapeutic schedules that manage chemo-resistance, antibiotic scheduling to avoid and even reverse antibiotic resistance in microbial populations, or the introduction of economic incentive packages to guide behavior). One of the more compelling potential applications of the methods developed in this paper is to frame people’s attitudes towards vaccination acceptance as a social contract [39, 40] and to devise dynamic incentive methods to encourage vaccination acceptance as well as to explore their theoretical limitations. Our method uses the Pontryagin maximum/minimum principle along with the 2 2 replicator dynamical system, with contraints, to determine schedules over one cycle time *T*, then we extend the results adaptively over multiple cycles *nT*. We show this leads to the identification of an absolute maximizer and minimizer 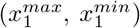 for the aggressor population, both of which are functions of the cycle time *T*. We believe the general framework layed out in the paper can be extended to *N N* replicator systems, as well as discrete (stochastic) models for the interaction of a finite number of partcipants using a Moran process, and we are currently extending the methods in this paper to include those settings.

## ACKNOWLEDGMENTS

We gratefully acknowledge support from the Army Research Office MURI Award #W911NF1910269 (2019-2024).

## Notes

### Competing Interest Statement

The authors have declared no competing interest.

